# Genomic analysis of worldwide sheep breeds reveals *PDGFD* a major target of fat-tail selection in sheep

**DOI:** 10.1101/2020.05.08.085431

**Authors:** Kunzhe Dong, Min Yang, Jiangang Han, Qing Ma, Jilong Han, Ziyi Song, Cuicheng Luosang, Neena Amatya Gorkhali, Bohui Yang, Xiaohong He, Yuehui Ma, Lin Jiang

**Affiliations:** Institute of Animal Sciences, Chinese Academy of Agricultural Sciences (CAAS), No. 2 Yuanmingyuan West Road, Beijing 100193, China; Key Laboratory of Animal (Poultry) Genetics Breeding and Reproduction, Ministry of Agriculture and Rural Affairs, CAAS, Beijing 100193, China; College of Animal Science and Technology, Shihezi University, Shihezi 832000, China; Research Center of Grass and Livestock, Ningxia Academy of Agriculture and Forestry Sciences, Yinchuan 750002, China; College of Animal Science and Technology, Guangxi University, Nanning, Guangxi 530004, China; Research Institute of Animal Science, Tibet Academy of Agricultural and Animal Husbandry Sciences, Lhasa 850000, China; Lanzhou Institute of Husbandry and Pharmaceutical Sciences, Chinese Academy of Agricultural Sciences (CAAS), Lanzhou 730050, China

**Keywords:** *PDGFD*, fat-tailed sheep, fat deposit, genomic scan

## Abstract

Fat tail is a special trait in sheep acquired during sheep domestication. Several genomic analyses have been conducted in sheep breeds from limited geographic origins to identify the genetic factors underlying this trait. Nevertheless, these studies obtained different candidates. The results of these regional studies were easily biased by the breed structures. To minimize the bias and distinguish the true candidates, we used an extended data set of 968 sheep representing 18 fat-tailed breeds and 14 thin-tailed breeds from around the world, and integrated two statistic tests to detect selection signatures, including Genetic Fixation Index (*F_ST_*) and difference of derived allele frequency (ΔDAF). The results showed that *platelet derived growth factor D (PDGFD)* exhibited the highest genetic differentiation between fat- and thin-tailed sheep breeds. Further analysis of sequence variation identified that a 6.8-kb region within the first intron of *PDGFD* is likely the target of positive selection and contains regulatory mutation(s) in fat-tailed sheep. Histological analysis and gene expression analysis demonstrated that *PDGFD* expression is associated with maturation and hemostasis of adipocytes. Luciferase reporter assays showed that a segment of conserved sequence surrounding the orthologous site of one sheep mutation is functional in regulating *PDGFD* expression in human. These results reveal that *PDGFD* is the predominant factor for the fat tail phenotype in sheep by contributing to adiopogenesis and maintaining the hemostasis of mature adipocytes. This study provides insights into the evolution of fat-tailed sheep and has important application to animal breeding, as well as obesity-related human diseases.

## Introduction

Following domestication in the Fertile Crescent approximately 8,000-9,000 years ago [1], sheep has spread and distributed over a wide geographical range worldwide mainly due to their adaptability to nutrient-poor diets and extreme environments, and thus developed a substantial variation in many phenotypic traits [2]. One of the main morphological changes is the lengthening of the tail and distinct patterns of tail fat deposition. It is believed that fat-tailed sheep was selected in response to the steppe and desert conditions in central Asia from 3, 000 BC [3], which is several thousand years after the domestication of its thin-tailed ancestor, and then spread east into north China and west into South Africa.

However, nowadays, fat-tail phenotype loses its earlier advantages and a decrease in the size of sheep tail is desirable for both producers and consumers, partially because that fat tails have a significant influence on fat deposition in other parts of the body [4], mating and normal locomotion of the animal [5]. Besides, the consumers in many instances show an increasing preference for lean meat. Genetic improvement is a more attractive strategy to develop sheep with small size of tail than the traditional practices in farms, like tail docking, as it is reliable, long lasting and humanism. For this purpose, finding the genes underlying the fat tail phenotype is the first and most important thing. Furthermore, elucidating the physiology of fat deposition in sheep tail provides significant insight into human health problems, such as obesity that is a major threat to the quality of human life in modern society [6].

Several efforts have been made aiming to hunt for genes or genomic regions associated with fat tail phenotype by genome-wide scans [7–12]. Nevertheless, the results of these studies remain controversial, with almost no consensus on their implications. This is quite surprising, as all the fat-tailed sheep seem to be of same origin with relatively short history and thus they are expected to have the similar genetic basis underlying the trait. Additionally, these previous studies suffered several limitations that hinder the search for valid candidates. First, they investigated sheep populations from relatively limited areas, which may identify the wrong candidate loci that were selected by other confounding factors such as geographic isolation. Second, these studies mostly applied allele frequency-based methods, such as Genetic Fixation Index (*F_ST_*) method, to identify candidate genes. *F_ST_*-based approach is widely used to identify the region with high divergence between populations; however, it lacks the power of telling the direction of selection. To clarify the controversy, here we performed a comprehensive genomic analysis of 18 thin-tailed and 14 fat-tailed sheep breeds from around the world, by integrating two selection tests, including Genetic Fixation Index (*F_ST_*) and the difference of derived allele frequency (ΔDAF) (referred to as DAF_Fat-tailed sheep_ – DAF_Thin-tailed sheep_), to identify positively selected genes specifically in fat-tailed sheep. This analysis revealed *PDGFD* as the major candidate underlying fat tail phenotype in sheep. Further histological and gene expression analysis demonstrated that *PDGFD* expression is associated with adipogenesis in fat tissues during sheep embryonic development and remains higher in fat tissues of fat-tailed sheep than that of thin-tailed sheep at both embryonic and adult stage. Finally, we carried out dual luciferase reporter assays and found that a segment of conserved sequence surrounding the orthologous site of one sheep mutation is functional in regulating *PDGFD* expression in human.

## Materials and methods

### Samples and genotyping

Genome-wide SNP data of five South Asian thin-tailed sheep breeds (*i.e*., Tibetan, Changthangi, Deccani, IndianGarole and Garut sheep), six European thin-tailed sheep breeds (*i.e*., Churra, Leccese, Comisana, Altamurana, MacarthurMerino and MilkLacaune sheep), three American thin-tailed sheep breeds (*i.e*., BarbadosBlackBelly, MoradaNova and SantaInes sheep), six Middle East fat-tailed sheep breeds (*i.e*., Afshari, LocalAwassi, Karakas, Norduz, Moghani and CyprusFatTail sheep), four African fat-tailed sheep breeds (*i.e*., EthiopianMenz, NamaquaAfrikaner, RedMaasai and RonderibAfrikaner) and eight Mouflon sheep individuals (wild sheep) were downloaded from Ovine HapMap project (http://www.sheephapmap.org/hapmap.php) [13]. SNP data of four Chinese fat-tailed sheep breeds (*i.e*., Hu, Tong, Large tail Han and Lop sheep) were obtained from a previous study [10]. SNP data of four Nepalese thin-tailed sheep breeds (*i.e*., Bhyanglung, Baruwal, Lampuchhre and Kage sheep) were generated in our lab [14]. Tail type for these breeds was determined by either direct observation or description of previous literatures. The detailed information of these sheep breeds was provided in **Supplementary Table S1**. All the SNP data were generated using the Illumina Ovine 50K Beadchip and were thus readily merged together. The final dataset included 968 sheep individuals and 47,415 common autosomal SNPs (based on genome Oar_v3.1).

### Determination of ancestral allele and data quality control

Ancestral allele information for a subset of 33,059 SNPs were obtained from Ovine HapMap [13]. For the remaining SNPs, the ancestral allele was deduced according to the major allele in Mouflon sheep. Finally, a total of 46,540 SNPs with available ancestral allele information were kept for next analysis. PLINK v2.05 [15] was applied for further SNP data quality control. SNPs were removed if any of the following conditions were met: 1) with call rate ≤90%; 2) with minor allele frequency (MAF) ≤0.05. Sheep individuals with an average call rate below 90% were discarded. To ensure independence among the studied sheep individuals, cryptic relatedness among individuals within each breed were identified using pair-wise Identity-By-Descent (IBD) metric (referred to as PI-HAT in PLINK). One individual from a pair of sheep individuals was removed from the following analyses if their PI-HAT value was over 0.3.

### Phylogenetic analysis

A pruned set of 32,450 SNPs were used to investigate the genetic relationship among these sheep breeds from different geographic locations. Principle Component Analysis (PCA) was performed with the GCTA software [16] and the individuals outside of their expected population clusters were excluded from further analysis. The neighbor-joining tree was constructed using PHYLIP 3.68 (http://evolution.genetics.washington.edu/phylip.html) on the basis of allele frequency data. After PCA analysis, a total of 45,337 SNPs for 828 individuals from 30 diverse sheep breeds were retained for downstream analyses (**Supplementary Table S1**).

### Genomic screen for positively selected genes in fat-tailed sheep

Three previous studies compared the genomic variations between Middle East/European thin-tailed versus (v.s.) Middle East fat-tailed sheep [11], European thin-tailed v.s. Middle East fat-tailed sheep [12], and Chinese thin-versus fat-tailed sheep [10], respectively. Therefore, we used sheep breeds from South Asia, Middle East and Europe to identify the positively selected genes in fat-tailed sheep. Three group-pair comparisons between thin- and fat-tailed sheep were considered, including Middle East fat-tailed sheep (MEF) v.s. South Asian thin-tailed sheep (SAT), MEF v.s. European thin-tailed sheep (EUT), and Chinese fat-tailed sheep (CHF) v.s. SAT.

Two statistics, including the *F_ST_* and ΔDAF were applied to evaluate the genetic differentiation of each SNP between thin- and fat-tailed sheep populations. The *F_ST_* analysis was conducted using Genepop 4.3 software [17]. ΔDAF was calculated as the derived allele frequency in the fat-tailed sheep minus the DAF in thin-tailed sheep (DAF_Fat-tailed sheep_ – DAF_Thin-tailed sheep_) using R version 3.3.3 (https://www.r-project.org/). For each group-pair comparison, *F_ST_* and ΔDAF at each SNP marker was calculated between each thin-tailed breed against each fat-tailed breed and averaged across breed-pairs to produce an overall value for each SNP. The top 1% SNPs with large overall *F_ST_* or ΔDAF value were considered as the significant SNPs in each test and the significant SNPs overlapped in both tests were considered as positively selected SNPs in fat-tailed sheep. Finally, positively selected SNPs that were identified in all three group-pair comparisons were considered as the loci under positive selection in fat-tailed sheep and genes within 150 kb of these loci were retrieved from Ensembl BioMart database (http://useast.ensembl.org/biomart/martview/625a397ecca62a421509935f099cdaa7).

Because the high average value of *F_ST_* or ΔDAF may be resulted from the extremely large values in several specific breed-pairs due to population structure, we additionally examined the DAF of the candidate SNPs in three thin-tailed sheep breeds from Americas and three fat-tailed sheep breeds from Africa and further filtered the promising candidate list by removing SNPs with DAF >0.5 in more than five thin-tailed sheep breeds or DAF <0.5 in more than four fat-tailed sheep breeds (>30% of total thin- or fat-tailed sheep breeds).

### Analysis of sequence variations within top genes

Genomic variations stored in Variant Call Format (VCF) file for 16 thin-tailed sheep individuals and 13 fat-tailed sheep individuals from 17 different breeds from around the world were downloaded from NextGen of Ensembl Projects (http://projects.ensembl.org/nextgen/) (**Supplementary Table S2**). The VCF file was converted to PLINK PED file using VCFtools [18]. PLINK [15] was used to perform the SNP quality control (removing SNPs with call rate ≤90% or MAF ≤0.05) and to calculate the allele frequency of each SNP in thin- and fat-tailed sheep group. The absolute difference of allele frequency (ΔAF) of each SNP between fat - and thin-tailed sheep group were calculated using R.

### Validation of top SNPs in expanded samples using Sequenom MassARRAY

A total of 13 intronic SNPs of *PDGFD* gene with the highest ΔAF value (>0.8) and derived allele frequency larger than 0.8 in fat-tailed sheep were selected for further validation in an expanded cohort containing 200 Tibetan sheep (Thin-tailed sheep) 184 Tan sheep (fat-tailed sheep). The genotyping of these 13 SNPs was carried out with Sequenom MassArray system. Briefly, primers for PCR and for locus-specific single-base extension were designed with MassArray Assay Design 4.0. The PCR products were applied for locus-specific single-base extension reactions. The final products were desalted and transferred to a matrix chip. The resultant mass spectrograms and genotypes were analyzed with MassArray Typer software.

### Sections, Hematoxylin and Eosin (HE) staining, Oil Red O staining

Embryonic tail tissues were collected from Tan sheep (fat-tailed sheep) at four different embryonic stages including embryonic day 60 (E60), E70, E80 and E90, each containing four samples. Tissues were fixed with 4% paraformaldehyde over-night at 4°C and then embedded in paraffin. Sections were cut at 8-um thickness. For HE staining, sections were deparaffinized with two changes of Xylene (10 minutes each) and were rehydrated with 100%, 95% and 80% ethanol rinses (5 minutes each). After a brief wash in distilled water, sections were stained in hematoxylin solution for 5 minutes, washed in running tap water for 5 minutes, differentiated in 1% acid alcohol for 30 seconds and washed again with running tap water for 1 minute. Following bluing in 0.2% ammonia water for 30 seconds and a wash in running tap water for 5 minutes, sections were rinsed in 95%alcohol and stained in eosin-phloxine solution for 30 seconds. Sections were then dehydrated with a series of rinses in 80% alcohol (5 seconds), 95% alcohol (twice, 10 seconds each), 100% alcohol (twice, 5 minutes each), and then mounted. For Oil Red O staining, 0.5 g Oil Red O powder (Sigma) was evenly dissolved in 100 mL of isopropanol to prepare stock solution, which was then diluted with distilled water at the ratio of 3:2 and filtrated to make working solution. The sections were rinsed with 60% isopropanol and then stained with Oil Red O working solution for 15 minutes. Sections were then mounted and imaged for evaluation of droplet formation.

### RNA-sequencing and bioinformatics analysis

Fat tissues of tail from Tan sheep (fat-tailed) and Suffolk sheep (thin-tailed sheep) at three different developmental stages including E60, E70, E80 were collected. Total RNA was extracted by RNeasy Lipid Tissue Mini Kit (Qiagen) and the RNA-seq library was constructed with Dynabeads mRNA DIRECT Kit (invitrogen)for whole transcriptome RNA-seq analysis. Pair-end RNA-sequencing was performed on X-ten system (Illumina) in 150-bp length.

After removing adaptor sequence and low-quality reads, pass-filtered reads were then mapped to Ensembl sheep reference genome Oar_v4.0 using Tophat 2.1.1 [19]. The genes annotated in Ensembl were quantified with Cufflinks [20]. FPKM (Fragments Per Kilobase of exon per Million fragments mapped) were calculated from raw counts and used to quantify relative gene expression for each sample.

### Quantitative reverse transcription-PCR (qRT-PCR) analysis

Tail tissues of Tan sheep (fat-tailed) at different developmental stages including E60, E70, E80, E90 and adult stage (about 2 years old, female), as well as from Suffolk sheep (thin-tailed sheep) at E70 and adult stage (Bhyanglung) (about 2 years old, female) were collected. Total RNA from tail tissues were isolated with RNeasy Lipid Tissue Mini Kit (Qiagen). 0.8μg of Total RNA was utilized as template for RT with random hexamer primers using PrimeScript RT reagent Kit (Takara). qRT-PCR was performed with respective gene-specific primers (*PDGFD:* Forward: GGGAGTCAGTCACAAGCTCT, Reverse: AGTGGGGTCCGTTACTGATG; *ACTB*: Forward:TCTGGCACCACACCTTCTAC; Reverse: TCTTCTCACGGTTGGCCTTG). All samples were amplified in triplicate and all experiments were repeated at least 3 independent times. Relative gene expression was converted using the 2^-ΔΔCT^ method against the internal control house-keeping gene ACTB where ΔΔCT= (CT_experimental gene_ - CT_experimental ACTB_) - (CT_control gene_ - CT_control ACTB_). The relative gene expression in control group was set to 1.

### *PDGFD* expression in adipose tissues/cells in mouse and human

Micro-array data of different cell types isolated from human white adipose tissues by fluorescence-activated cell sorting (FACS) including CD45-/CD34+/CD31- (progenitors), CD45+/CD14+/CD206+ (total macrophages), CD45+/CD14+CD206+/CD11c+ (M1 macrophages), CD45+/CD14+/CD206+/CD11c- (M2 macrophages), CD45+/CD3+ (Total T cells), CD45+/CD3+/CD4+/CD8- (Th T-cells), CD45+/CD3+/CD4-/CD8+ (Tc T-cells) (GSE100795) [21] were obtained from GEO database and re-analyzed with online build-in tool GEO2R. Raw gene count matrix generated from RNA-seq for mouse 3T3-L1 preadiopocytes and differentiating cells at 24 hours after induction by DHA differentiation cocktail was obtained from GEO database (GSE118471) [22] and FPKM was used for quantification of gene expression level. Micro-array data of inguinal fat tissues from mice exhibiting high or low weight gain after 4 weeks on a high saturated fat diet (GSE4692) [23], adipocytes from non-diabetic lean and non-diabetic obese Pima Indian subjects (GSE2508) [24] were obtained from GEO database and analyzed with online build-in tool GEO2R.

### Dual luciferase reporter assay

Using human genomic DNA as template, two fragments (WT, 222 bp; Del, 202 bp) spanning *PDGFD* gene promoter region were synthetized directly by BerryGenomics Company (Beijing, China), and then the products were cloned into the site between Kpn1 and EcoRV restriction sites of pGL4.23 (Promega) luciferase reporter vector. All plasmids were sequenced to verify the integrity of the insert. Transfection was performed with Lipofectamine 3000 reagent (ThermoFisher) essentially following manufactory’s instruction. The promoter activity was evaluated by measurement of the firefly luciferase activity relative to the internal control minimal TK promoter driven-Renilla luciferas eactivity using the Dual Luciferase Assay System as described by the manufacturer (Promega). A minimum of four independent transfections was performed and all assays were replicated at least twice.

## Results

### Identification of genes underlying fat tail in sheep

SNP Beadchip data for a total of 968 sheep individuals was collected and used in this study. These samples contained eight wild Mouflon sheep individuals, 18 thin-tailed sheep breeds and 14 representative fat-tailed sheep breeds from different countries or regions within countries (**Fig. 1A; Supplemental Table S1**). The geographic distributions of the sheep breeds included in this report include South Asia, East Asia, Middle East, Europe, Africa and Americas (**Fig. 1A; Supplementary Table S1**). After a series of quality control filters, a total of 45,337 SNPs and 828 individuals from 30 diverse sheep breeds worldwide were kept for further analysis.

**Figure 1.**
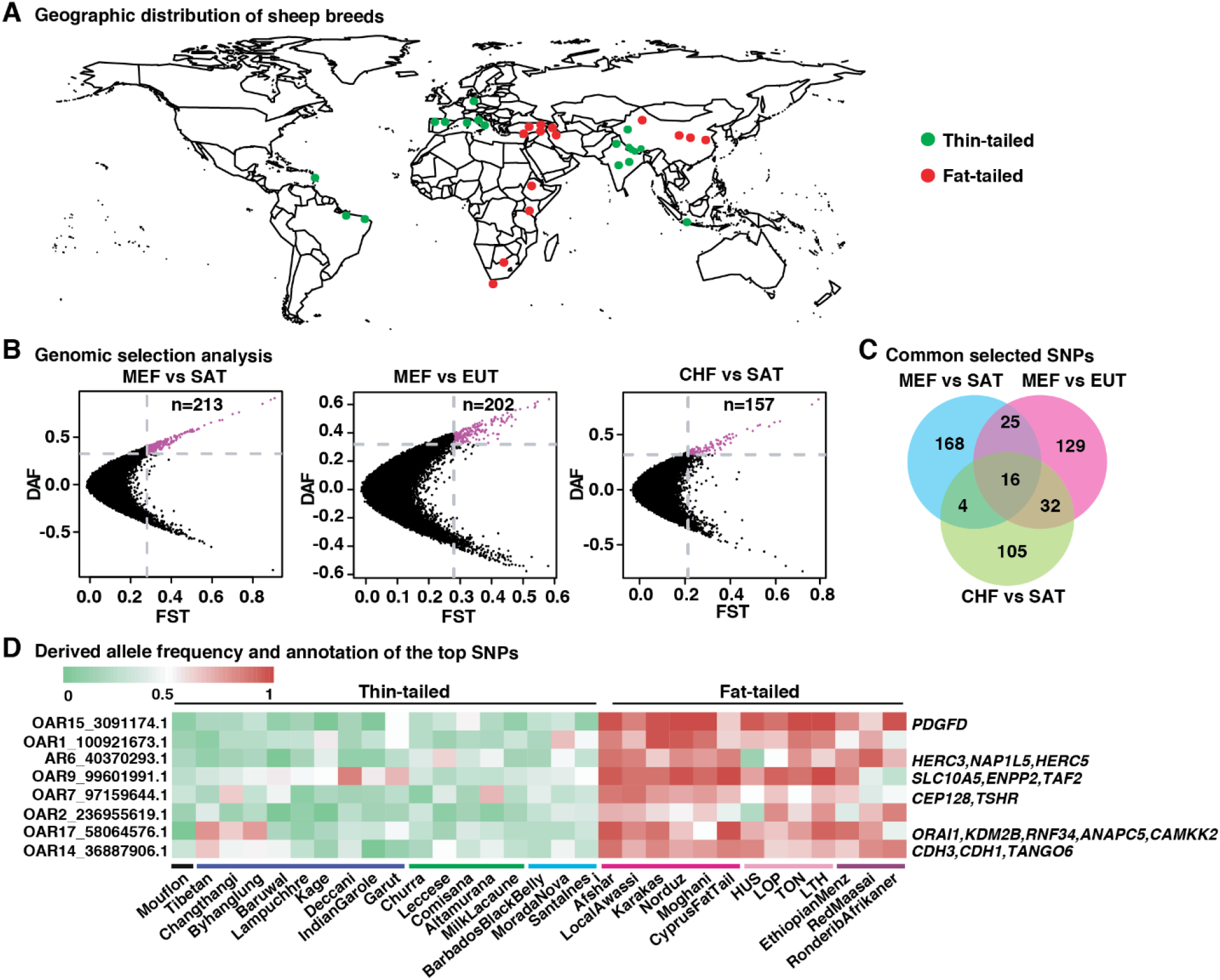
Identification of positively selected loci in fat-tailed sheep. **(A)** The geographic distribution of the sheep breeds used in this study, each of which is represented by a dot on the world map. **(B)** The number of positively selected loci identified in three different group-pair comparisons. MEF: Middle East fat-tailed sheep from Middle East; SAT: South Asian thin-tailed sheep; EUT: European thin-tailed sheep; CHF: Chinese fat-tailed sheep. **(C)** The number of positively selected loci identified in all the three group-pair comparisons. **(D)** The derived allele frequency (DAF) of the 16 positively selected loci identified in all three group-pair comparisons in all the studied sheep breeds. SNPs were sorted according to the average value of *F_ST_* and ΔDAF among the three group-pair comparisons indicated in Figure 1b from high to low. The important annotated genes of each SNP were labeled in the right.

Because the wild ancestor of domestic sheep, the Mouflon sheep, is thin-tailed, accordingly modern thin-tailed sheep is expected to keep initial allele state while fat-tailed sheep exhibits derived alleles at the loci related to fat tail phenotype. Under this scenario, we integrated two statistical measures, the *F_ST_* and the ΔDAF, to increase the power of identifying genomic loci specifically selected in fat-tailed sheep breeds. For comparison with previous studies [7–12], we separately performed *F_ST_* and ΔDAF analysis in three group-pair comparisons, including MEF vs SAT, MEF vs EUT, and CHF vs SAT. Totally, 213, 202 and 157 positively selected SNPs were identified by both *F_ST_* and ΔDAF test in the three comparisons, respectively (**Fig. 1b**) (See **Supplementary Tables S3-S5** for the full significant SNP list). Among them, 16 loci were common to all the three group-pair comparisons (**Fig. 1c**).

Eight out of the 16 common significant SNPs were further ruled out from our candidate list because they have DAF > 0.5 (or DAF < 0.5) in more than 30% of the thin-tailed (or fat-tailed) sheep breeds, after examining the DAF of these SNPs in additional three American thin-tailed sheep breeds and three African fat-tailed sheep breeds (See Methods) (**Supplementary Fig. S1**). The remaining eight SNPs are highly diverged between thin- and fat-tailed sheep, and keep ancestral allele in most thin-tailed sheep breeds while derived allele in most fat-tailed sheep breeds worldwide (**Fig. 1d**), indicative of promising candidates associated with fat tail phenotype. PCA and phylogenetic tree analysis using these eight SNPs showed clearly separated clades between thin- and fat-tailed sheep, which is quite distinct from the results obtained based on genome-wide variants showing geographic clustering (**Supplementary Fig. S2**) and further supported the strong divergence of these loci. Gene annotation of 150 kb-long genomic region surrounding the eight SNPs revealed 24 genes and *PDGFD* corresponding SNP OAR15_3091174 among them that is located on chromosome 15 was most highly differentiated (**Fig. 1d**).

### Analysis of sequence variations of *PDGFD* region

To better investigate these selected genes, we extracted their genome sequence variants of 16 thin-tailed and 13 fat-tailed sheep individuals with diverse geographic origins from NextGen project (**Supplementary Table S2**). A total of 17,124 SNPs were identified in the selected regions among these sheep individuals (MAF >0.05). We calculated the ΔAF of each SNP and confirmed that the *PDGFD* gene is the most divergent locus between fat- and thin-tailed sheep (**Fig. 2A**), again implying that *PDGFD* is likely the most promising candidate for the sheep tail divergence. PDGFD is one of the platelet-derived growth factor (PDGF) family members, which play an important role in regulation of adipocyte development and function [25, 26]. The expression pattern across multiple issues obtained from Human Protein Atlas database (http://www.proteinatlas.org/) showed that *PDGFD* is relatively more abundantly expressed in adipose tissue than that in other tissues (**Supplementary Fig. S3**). These already known involvements of *PDGFD* in lipid metabolism provide further support for the hypothesis that this gene is an ideal candidate for fat tail in sheep.

**Figure 2.**
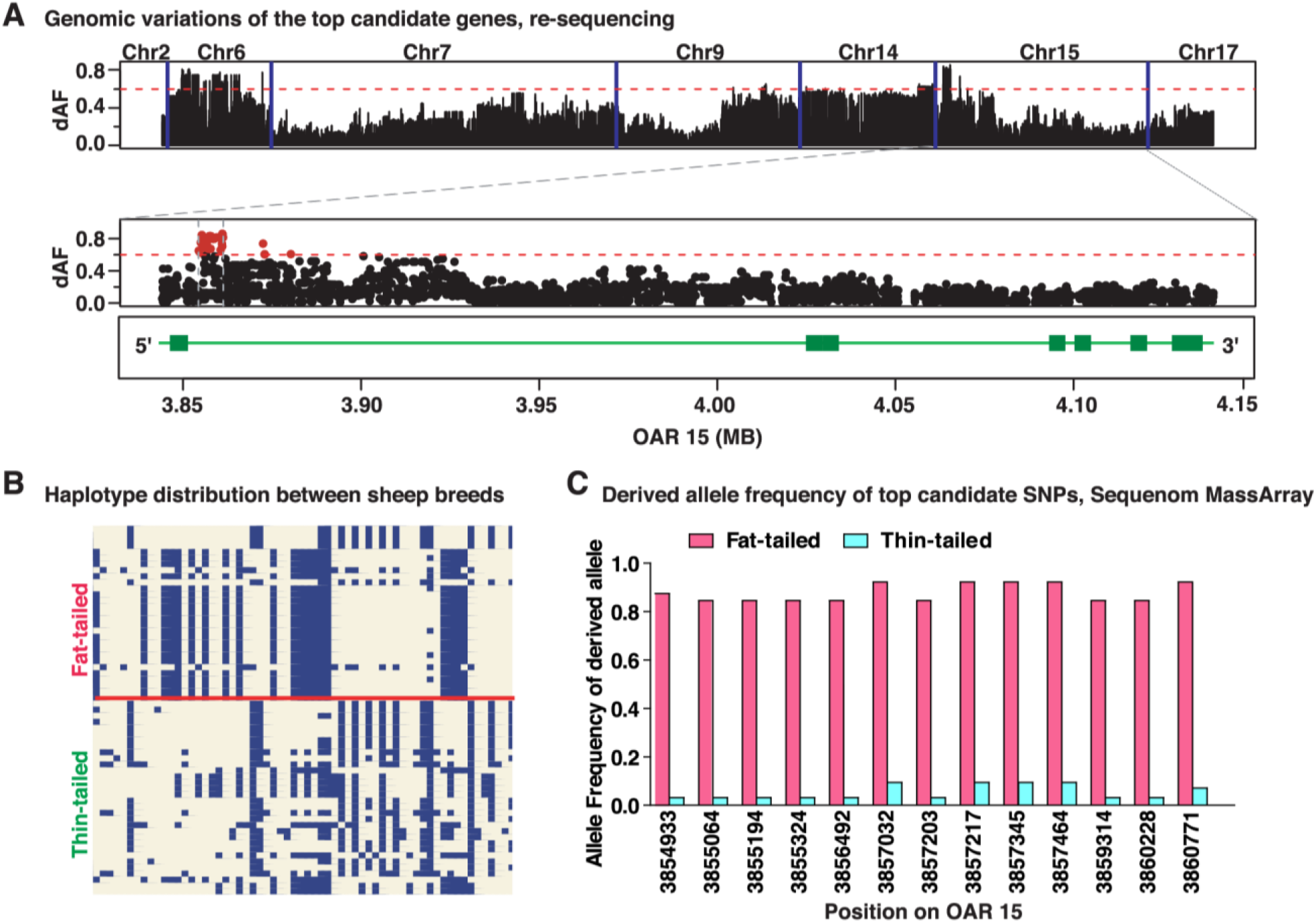
Analysis of sequence variations of *PDGFD* gene. **(A)** The distribution of the absolute difference of allele frequency ΔAF of sequence variation within the 24 identified genes with selection signatures between 13 fat-tailed sheep individuals and 16 thin-tailed sheep individuals from different regions. The sequence variations were downloaded from NextGene project. The red dashed line indicates the threshold value (ΔAF >0.6). The bottom panel shows the distribution of the absolute difference of allele frequency ΔAF of sequence variation within *PDGFD* gene. The red dashed line indicates the threshold value (ΔAF >0.6) and red dots represents SNPs with ΔAF >0.9. **(B)** Haplotype compassion between thin-tailed and fat-tailed sheep obtained based on all the variations within the highly differentiated 6.8-kb genomic interval (Chr15:3,854,063-3,860,894 bp). Each column is a polymorphic genomic location (122 in total), each row is a phased haplotype (16 thin-tailed sheep and 13 fat-tailed sheep individuals), and alternative alleles are labelled in blue. **(C)** Derived allele frequency of the 13 top candidate SNPs in additional fat-tailed sheep (Chinese Tan sheep) and thin-tailed sheep (Chinese Tibetan sheep) obtained by Sequenom MassARRAY.

Furthermore, ΔAF is noted to be particularly elevated in a 6.8-kb region (Chr15: 3,854,063 - 3,860,894 bp) in *PDGFD* gene containing 51 most differentiated SNPs (ΔAF >0.6) (**Supplementary Table S6**). Haplotype comparison analysis using all the SNPs (n=122) within this region revealed a consistent differentiation between sheep breeds with different tail types (**Fig. 2B**). Therefore, it is likely that this 6.8-kb region is the target of positive selection for fat tail and is the best candidate region for the functional mutation(s). This region is located at the first intron of *PDGFD* (**Fig. 2B**, bottom), suggesting that the mutations causing the association with the fat-tail phenotype are regulatory. We next genotyped a total of 13 SNPs that exhibit highest ΔAF value (>0.8) between thin- and fat-tailed sheep and large derived allele frequency in fat-tailed sheep (>0.8) based on genome re-sequencing results, using Sequenom MassARRAY in an expanded cohort containing 200 Tibetan sheep (Thin-tailed sheep) 184 Tan sheep (fat-tailed sheep). This analysis confirmed that all these SNPs are highly divergent (**Fig. 2C**). Collectively, these observations strongly suggested that the 6.8-kb region identified here deserves further investigation for revealing the causative mutation(s).

In addition to *PDGFD*, several other selected genes identified in this study have known functions that are associated with lipid metabolism, such as *ENPP2* [27], *ANAPC5* [28], *RNF34* [29], *KDM2B* [30], *CAMKK2* [31], *TSHR* [32], and *CDH1* [33]. The genetic differentiation for these genes was not as pronounced as for the *PDGFD* loci, implying that these loci likely contribute to the phenotypic difference with relatively small effects. We also examined the DAF distribution of the top candidate SNPs that were proposed in previous studies and the results showed that the majority of these SNPs are not consistently diverged between thin- and fat-tailed sheep populations [7–12] (**Supplemental Fig. S4**).

### *PDGFD* expression is associated with adipogenesis

As an initial step to elucidate the role of *PDGFD* in the development of fat tail in sheep, tail tissues from a fat-tailed Chinese sheep breed, namely Tan sheep, at four different embryonic time points, including embryonic day 60 (E60), E70, E80 and E90, were collected and subjected to histological analysis. HE staining showed that lipid vacuole is absent in cells at E60 and E70, while accumulates in cells at E80 and E90, as evidenced by observation that one large vacuole was present in the cell and the nucleus was packed in a corner (**Fig. 3A**), a typical feature of mature adipocytes. In line with this, Oil Red O staining which visualizes fat-containing droplets revealed that a large portion of cells at E80 and E90 showed distribution of lipid drops (**Fig. 3B**). These observations suggestedthat committed preadioocytes are present in tail tissues at E60 and E70 which differentiate to mature adipocytes around E80 and E90.

**Figure 3.**
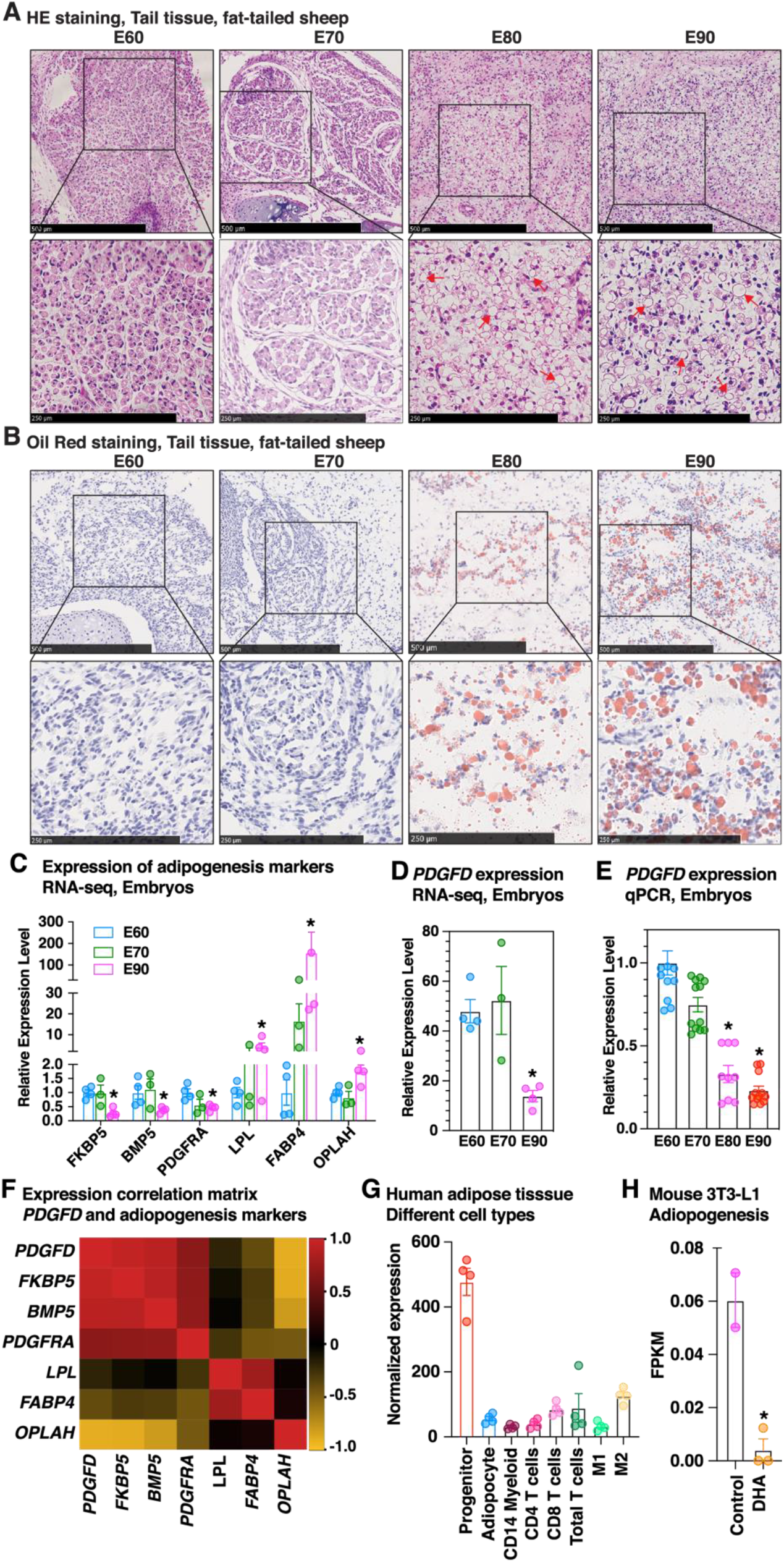
*PDGFD* expression is associated with adiopogenesis. **(A)** Hematoxylin and Eosin (HE) staining for tail tissues of fat-tailed sheep at embryonic day 60 (E60), E70, E80 and E90. The boxed areas are magnified on the bottom. Red arrows point to representative lipid drops. **(B)** Oil Red O staining for tail tissues of fat-tailed sheep at E60, E70, E80 and E90. The boxed areas are magnified on the bottom. **(C)** Expression of several marker genes involved in adipogenesis in tail tissues of fat-tailed sheep at different stages of embryonic development revealed by RNA-seq. *PDGFD* expression in tail tissues of fat-tailed sheep at different stages of embryonic development revealed by **(D)** RNA-seq and **(E)** qPCR. **(F)** Heatmap showing the correlation matrix between expression of *PDGFD* and the indicated adipogenesis markers. **(G)** Expression of *PDGFD* in different cell fractions of human subcutaneous adipose tissues. Macrophages were subdivided into M1 and M2 (GSE100795). **(H)** Expression of *Pdgfd* during the adipogenesis of mouse 3T3-L1 preadiopocytes induced by ω-3 fatty acid DHA revealed by RNA-seq (GSE118471).

Further RNA-seq analysis revealed that the expression of some marker genes which are known to be down-regulated *(FKBP5, BMP5* and *PDGFRA)* and up-regulated (*LPL*, *FABP4* and *OPLAH)* during adipogenesis had no obvious difference between E60 and E70 while significantly decreased and increased at E80 compared to earlier stages, respectively (**Fig. 3C**), confirming the histological results that cells in tail tissues of fat-tailed sheep undergo adipogenesis from E60/70 to E80/90. Interestingly, *PDGFD* expression was undistinguishable between E60 and E70, but was dramatically decreased at E80 (**Fig. 3D**). Subsequent qRT-PCR analysis confirmed the large reduction of *PDGFD* expression at E80, followed with a continued but not significant decline at E90, as compared to that at E60 and E70 (**Fig. 3E**). Other candidate genes including *BMP2* which were proposed by several studies [8, 9, 12], exhibited no specific oscillation in expression mirroring the process of adipocyte maturation in tail tissues along with embryonic development (**Supplemental Fig. S5**). Gene expression correlation analysis revealed that *PDGFD* expression is highly positively correlated with markers enriched in preadipocytes while reversely correlated with markers up-regulated in mature adipocytes (**Fig. 3F**). Accordantly, retrospective analysis of public transcriptomic data sets revealed that *PDGFD* expression is higher in adipose progenitors than that in other cell types isolated from human white adipose tissues (**Fig. 3G**) and is dramatically reduced during the adipogenesis of mouse 3T3-L1 (**Fig. 3H**). These results together suggest that *PDGFD* expression is enriched in preadipocytes and is down-regulated during adipogenesis.

### *PDGFD* is dysregulated between adipose tissues of lean and fat individuals across species

Further qRT-PCR analysis showed that *PDGFD* expression is higher in tail tissues of fat-tailed sheep than that in thin-tailed sheep at both E70 and adult stages (**Fig. 4A and B**). Interestingly, re-analysis of public data sets revealed that *PDGFD* is also more abundantly expressed in adipose tissues of obese individuals than that of lean individuals in both mouse (**Fig. 4C**) and human (**Fig. 4D and E**), suggesting an evolutionarily conserved role of *PDGFD* involving in hemostasis of mature adipocytes.

**Figure 4.**
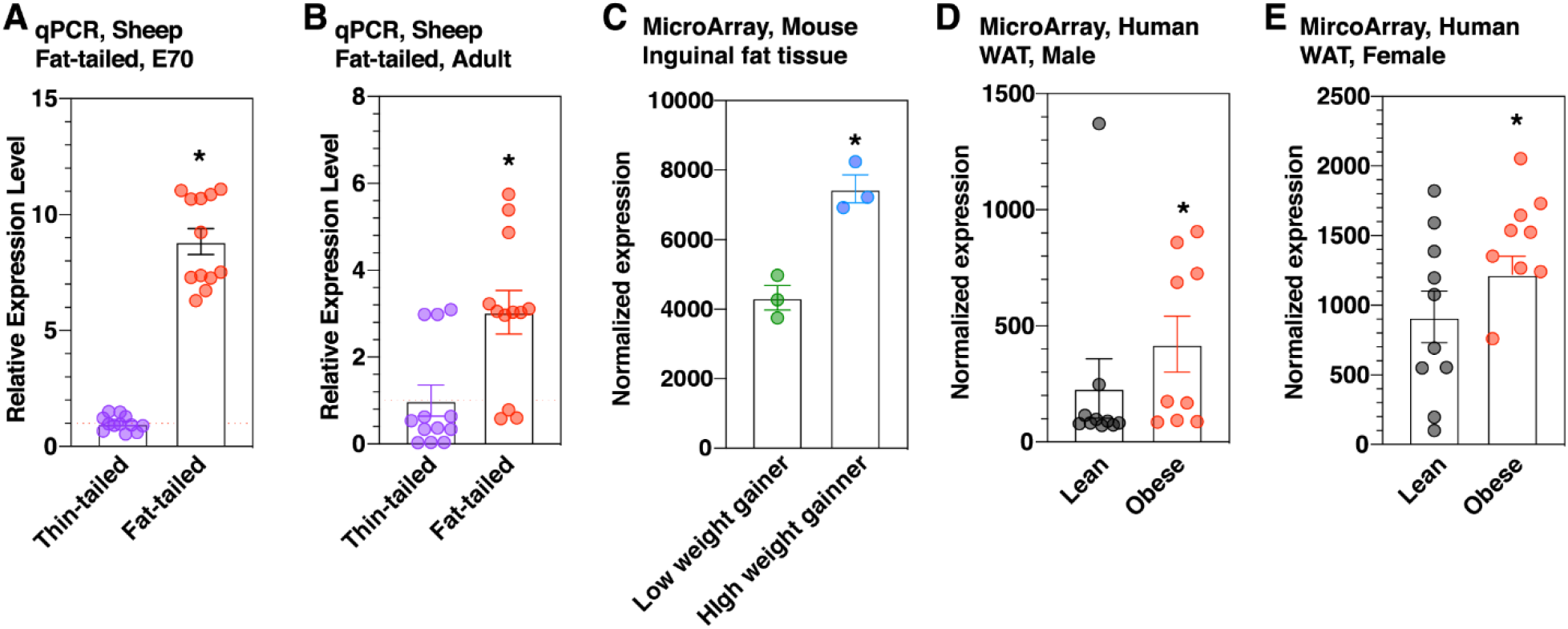
*PDGFD* expression is higher in adipose tissues of fat individuals than that of lean individuals across species. **(A)** *PDGFD* expression is higher in fat tissues of thin-tailed and fat-tailed sheep at E70 and **(B)** adult stage revealed by qPCR. *P <0.05. **(C)** *Pdgfd* expression is higher in inguinal fat of high weight gaining mice than that of in low weight gaining mice revealed by microArray analysis (GSE4692). *P <0.05 (n=3). *PDGFD* expression is higher in adipocytes from lean than that from obese Indian individuals in both **(D)** male and **(F)** female subjects revealed by microarray (GSE2508).

### A segment of conserved intronic region affects *PDGFD* expression in human

We next sought to explore the regulatory role of the most highly differentiated SNPs found within the first intron of *PDGFD* (**Fig. 2**). It was noted that a mutation (SNP OAR15_3859314) displays high frequency in fat-tailed sheep (**Fig. 5A**) and occurs at extremely conserved region between species including human (**Fig. 5B**). We therefore suspected that it could be critical for regulating *PDGFD* expression and selected it for further investigation. Together with the previous results showing that similar to observations in sheep, *PDGFD* expression is also enriched in human preadipocytes and higher in obese human individuals (**Fig. 3G and Fig. 4E-F**), we set out to extend our finding in sheep to human by investigating human *PDGFD* orthologous region. To do this, we cloned a fragment of human sequence (WT, wild type) as well as a mutated fragment with depletion of a conserved 20-bp region surrounding the site orthologous to sheep SNP OAR15_3859314 (Del) into a luciferase reporter vector and performed dual luciferase reporter assays (**Fig. 5C**). We first transduced the vectors into human A549 cells and the results showed that the activity of reporter carrying WT human sequence is remarkably suppressed as compared to blank luciferase vector (**Fig. 5D**), suggesting that the inserted WT sequence plays an inhibition role in regulating *PDGFD* expression. Interestingly, activity of the mutated reporter was restored relative to the reporter carrying the WT sequence (**Fig. 5D**), indicating that the conserved region is functional and is capable of transcriptionally activating the *PDGFD* expression. These observations were further confirmed by luciferase reporter assay performed in mouse 3T3-L1 preadipocytes (**Fig. 5E**). More interestingly, eQTL (Expression quantitative trait loci) analysis revealed that a mutation located 532 bp immediately downstream of the corresponding locus of sheep OAR15_3859314 mutation in human genome significantly increases the expression of *PDGFD* in human adipose tissues (**Fig. 5F**).

**Figure 5.**
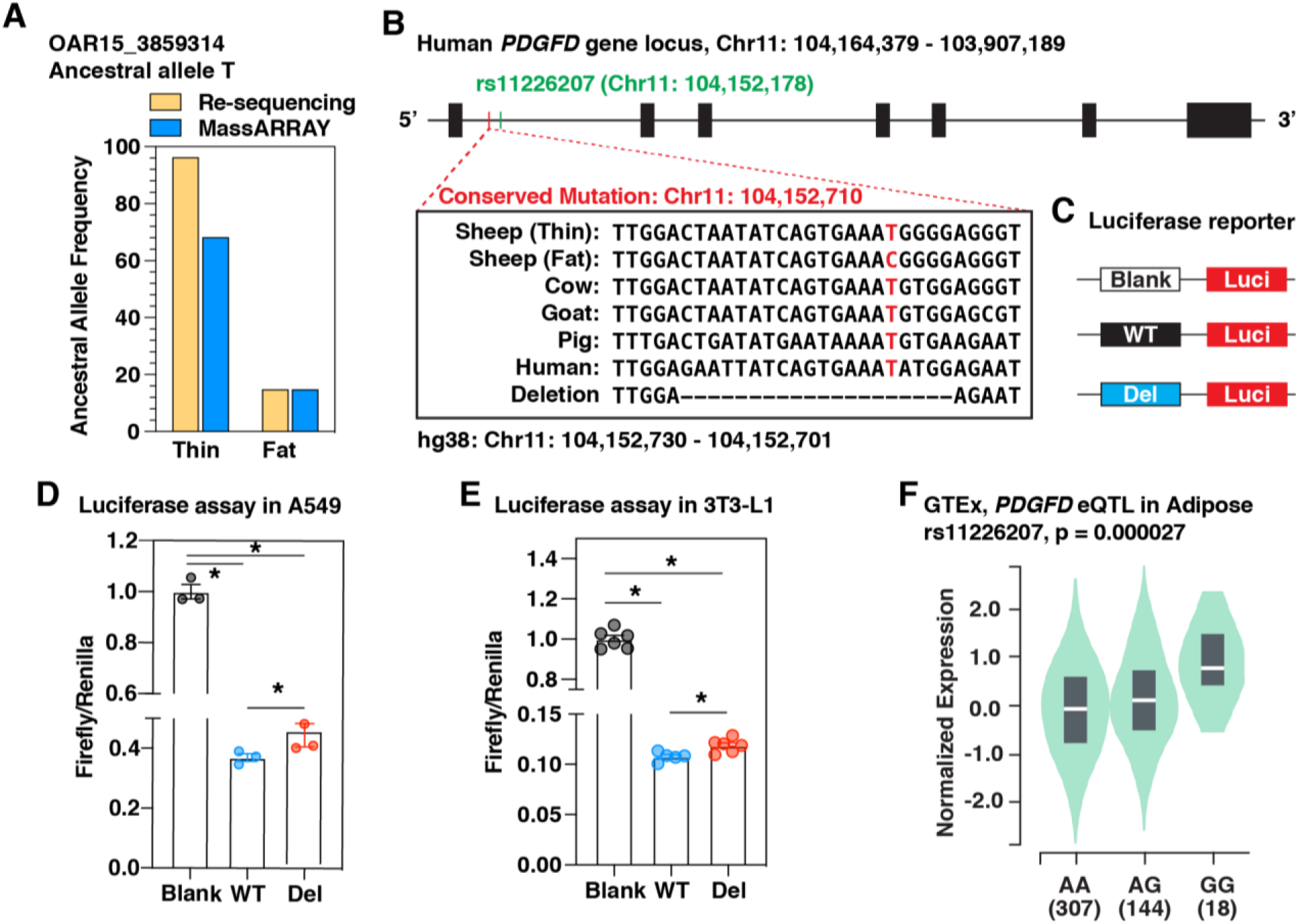
A mutation occurring at conserved site of *PDGFD* promoter regulates *PDGFD* expression. **(A)** Ancestral allele frequency of intronic SNP chr15:3,859,314 in thin- and fat-tailed sheep breeds determined by genome re-sequencing (in 29 sheep individuals) and massARRAY (in expanded samples). **(B)** Schematic diagram of the human *PDGFD* gene structure showing the site corresponding to sheep intronic SNP OAR15_3859314 (in red) is conserved across multiple species. The wild type sequence and sequence that deletes a 20-bp fragment containing the conserved mutation (deletion) were used for luciferase reporter construction. The locus in green (rs11226207) indicates the SNP described in “**F**”. **(C)** Schematic diagram showing the strategy of luciferase reporter construction. A 222-bp fragment within human first intron containing the homologous region of sheep intronic SNP chr15:3,859,314 and mutated version with depletion of 20-bp sequence harboring the conserved mutation were cloned into Pgl4.23 luciferase reporter vector. The blank vector served as control. **(D)** Results of luciferase reporter assays performed in A549 and (**E**) 3T3-L1 cells. Values of blank vector were set to 1. (**F**) A mutation proximal to SNP chr15:3,859,314 significantly alters *PDGFD* expression in human adipose tissues revealed by eQTL analysis in GTEx database. The location of this SNP is indicated in “**B**”.

## Discussion

To accurately map the candidate gene(s) underlying the fat tail phenotype of sheep, herein we comprehensively analyzed genomic variation data from a large cohort of sheep breeds with different tail types from around the world and integrated two different selection tests to detect genomic regions with signals of selection. We demonstrated that *PDGFD* gene loci exhibited the highest genetic differentiation between fat- and thin-tailed sheep and further found that the potential causal mutations are located with regulatory region. In addition, we show that *PDGFD* expression is negatively associated with maturation of adipocytes and is higher in fat tissues of fat individuals than that in lean individuals across different species. Finally, we provide evidence showing that a mutation occurring at conserved region of first intron is functional to transcriptionally activate the expression of *PDGFD* in human.

Compared to several existing papers that have studied sheep breeds from limited geographical areas and reported quite different candidates associated with fat tail phenotype [7–12], this study analyzed the largest cohort of samples until so far, to the best of our knowledge, originating from around the world. This practice avoids bias arising from geographic isolation or population structure and thus represents a more unambiguous and powerful strategy. Actually, our replication analysis of top SNPs proposed by previous studies in our large cohort reveals that most of these SNPs show genetic differentiation only between some regional sheep populations including SNPs corresponding to *BMP2*, which have been reported by several independent studies [8, 9, 12]. For instance, three SNPs, OAR13_51852034.1, ORA13_51886803.1 and s27419.1, at the *BMP2* locus, exhibited relatively higher DAF in fat-tailed sheep than that in thin-tailed (**Supplemental Fig. S4**). Unfortunately, the genetic differentiation level of the former two SNPs is below the genome-wide threshold in one (MEF vs SAT) and two group-pair comparison(s) (MEF vs SAT, MEF vs EUT), respectively (**Supplemental Table S3-5**), which are probably caused by the low abundance of derived allele in some fat-tailed sheep populations, such as Afshari, LocalAwassi and EthiopianMenz (**Supplemental Fig. S4**). Therefore, *BMP2* has lower priority than the *PDGFD* locus as the candidate for fat-tailed phenotype. Furthermore, a recent study based on whole-genome re-sequencing data revealed that *PDGFD* is the most highly differentiated loci between fat-tail sheep populations from China and Middle East and thin-tailed Tibetan sheep, followed by *BMP2* as the second top highly divergent gene [9], providing further support to our finding that *PDGFD* is the predominant candidate for fat tail in sheep, although our study suffers limitation that only genes covered by the Ovine 50K Beadchip were investigated.

Adipose tissue development is associated with specification and differentiation of precursor cells (preadipocytes) to mature adipocytes as well as expansion of adipocyte size [34]. This finely orchestrated process involves changes in expression of a series of molecular regulators which remain to be discovered. Our histological analysis and gene expression analysis suggest that *PDGFD* expression is negatively associated with adipocyte accumulation in fat tissues during embryonic development in sheep (**Fig. 3A-F**). Consistently, re-analysis of data generated from previous studies in human and mouse reveals that *PDGFD* expression is enriched in preadipocytes and is decreased during the adipogenesis from precursor cells (**Fig. 3G-H**) [21, 22]. These observations strongly suggest that *PDGFD* plays a critical role in the initiation and/or commitment of preadipocytes and this function is conserved and universe across species. Despite of a continuous decline of expression during development, we find that *PDGFD* sustains higher expression in tail tissues of fat-tailed sheep than that of thin-tailed sheep from embryonic (E70) to adult stage (**Fig. 4A and B**). Supporting this, a previous study reported that the expression of *PDGFD* is significantly upregulated in tail adipose tissue from fat-tailed Hulun Buir sheep as compared to thin-tailed Tibetan sheep [35]. Furthermore, another study showed that *PDGFD* is differentially expressed between tail adipose tissues from female Hulun Buir sheep with different levels of tail fat accumulation [36]. More interestingly, this expression pattern holds true in mouse and human, with obese individuals exhibiting higher level of *PDGFD* expression in adipose tissues (**Fig. 4C-E**) [23, 24]. Together with a recent study reporting that *PDGFD* expression is associated with adipocyte number in human white adipose tissue [37], we propose that *PDGFD* is a novel regulator of adipogenesis and adipocyte hemostasis. Further gain/loss-of-function assays are necessary to confirm this potential function.

We discover a list of mutations that are located within the first intron of *PDGFD* and display high frequency in fat-tail sheep while low abundance in thin-tailed sheep (**Fig. 2 and Supplementary Table S6**). Since intronic mutations mainly affect the transcriptional efficiency of genes by creating or disrupting binding sites for transcriptional factors [38, 39], it is likely that the elevated expression of *PDGFD* in fat-tailed sheep is attributed to some/one of these mutations. We focus on one mutation that occurs within evolutionarily conserved region (**Fig. 5A-B**) and investigate the regulatory function of the conserved segment in human. Interestingly, we prove that a fragment of human orthologous sequence suppresses the expression of *PDGFD*, while removal of the surrounding sequence of this mutation attenuates the inhibition effect (**Figs. 4A-D**). It suggests that the conserved segment surrounding the mutation site (sheep SNP OAR15_3859314) transcriptionally activates the expression of *PDGFD* and explains the higher expression of *PDGFD* in tail tissues of fat-tailed sheep. Of note, this mutation is not present in human and other species, which mirrors the fact that fat tail phenotype is unique in sheep and also suggests a conserved role of the intronic region in regulating *PDGFD* expression. Interestingly, a mutation located 532 bp downstream and supposedly to be within the same linkage disequilibrium block of corresponding site of sheep SNP OAR15_3859314 significantly increase *PDGFD* expression in human adipose tissues (**Fig. 5F**). Coupled to the observations that *PDGFD* expression in adipose tissues of obese individuals is higher than that of lean individuals in both sheep and human (**Fig. 4**), we consider that mutations occurring at the intronic region orthologous to the candidate region that we identified in sheep (**Fig. 2**) are prone to elevate *PDGFD* expression and contribute to fat accumulation.

In conclusion, we have demonstrated that *PDGFD* gene is the predominant factor underlying the fat tail phenotype in sheep and is a novel regulator for adiopogenesis and maintaining hemostasis of mature adipocytes. Mutations occurring within the first intron of *PDGFD* is likely associated with *PDGFD* transcriptional activity and fat deposition. These findings are important for understanding the evolution of the fat tail in sheep and useful in genome-based animal improvement, as well as obesity-related diseases in human.

## Acknowledgements

The ovine SNP50 HapMap dataset used for the analyses of this study was provided by the International Sheep Genomics Consortium (ISGC) and obtained from www.sheephapmap.org in agreement with the ISGC Terms of Access. This study makes use of the sheep re-sequencing data generated by ISGC. The project was supported by the National Natural Science Foundation of China (U1603232, 31702092, 31802045), Ningxia Key Research and Development Project (2018ZDKJ0123), the Agricultural Science and Technology Innovation Program of China (ASTIP-IAS01), the earmarked fund for Modern Agro-industry Technology Research System (CARS-40-01). LJ was supported by the Elite Youth Program in Chinese Academy of Agricultural Sciences. Special Funds for Basic Research of Young Scholars of Shihezi University (CXRC201808, RCSX2018B10).

## Author contributions

K.Z.D., L.J. and Y.H.M. conceived and designed the study. K.Z.D., M.Y., J.L.H. and B.H.Y. contributed to determination of tail type of all the sheep population used in this study. N.G. collected the Nepalese sheep blood samples. M.Y., J.G.H., J.L.H. and Q.M. contributed to sample collection, DNA extraction, genotyping and qRT-PCR analysis. J.G.H. contributed to the histological analysis. K.Z.D. and Y.M. performed the data analysis. K.Z.D. wrote the manuscript. Y.H.M., L.J., X.H.H. and Z.Y.S edited the manuscript. All authors reviewed and approved the manuscript.

## References

1. Calatayud Giner, S., [The spread of agricultural knowledge and the leading role of elites in the origins of contemporary agriculture: Valencia, 1840-60]. Hist Agrar, 1999(17): p. 99–127.

2. Kijas, J.W., et al., A genome wide survey of SNP variation reveals the genetic structure of sheep breeds. PLoS One, 2009. 4(3): p. e4668.

3. Ryder, M.L., Sheep & man. Sheep & man., 1983.

4. Farid, M., et al., Protein requirements for maintenance of Barki desert sheep [Egypt]. Journal of Agricultural Sciences, Mnsoura, 1984.

5. Kridli, R. and S. Said, Libido testing and the effect of exposing sexually naive Awassi rams to estrous ewes on sexual performance. Small Ruminant Research, 1999. 32(2): p 149–152.

6. Collaborators, G.B. D.O., et al., Health Effects of Overweight and Obesity in 195 Countries over 25 Years. N Engl J Med, 2017. 377(1): p. 13–27.

7. Mwacharo, J.M., et al., Genomic footprints of dryland stress adaptation in Egyptian fat-tail sheep and their divergence from East African and western Asia cohorts. Sci Rep, 2017. 7(1): p. 17647.

8. Mastrangelo, S., et al., Genome-wide scan of fat-tail sheep identifies signals of selection for fat deposition and adaptation. Animal Production Science, 2019. 59(5): p. 835–848.

9. Pan, Z., et al., Rapid evolution of a retro-transposable hotspot of ovine genome underlies the alteration of BMP2 expression and development of fat tails. BMC Genomics, 2019. 20(1): p. 261.

10. Yuan, Z., et al., Selection signature analysis reveals genes associated with tail type in Chinese indigenous sheep. Anim Genet, 2017. 48(1): p. 55–66.

11. Moradi, M.H., et al., Genomic scan of selective sweeps in thin and fat tail sheep breeds for identifying of candidate regions associated with fat deposition. BMC Genet, 2012. 13: p. 10.

12. Moioli, B., F. Pilla, and E. Ciani, Signatures of selection identify loci associated with fat tail in sheep. J Anim Sci, 2015. 93(10): p. 4660–9.

13. Kijas, J.W., et al., Genome-wide analysis of the world’s sheep breeds reveals high levels of historic mixture and strong recent selection. PLoS Biol, 2012. 10(2): p. e1001258.

14. Gorkhali, N.A., et al., Genomic analysis identified a potential novel molecular mechanism for high-altitude adaptation in sheep at the Himalayas. Sci Rep, 2016. 6: p. 29963.

15. Purcell, S., et al., PLINK: a tool set for whole-genome association and population-based linkage analyses. Am J Hum Genet, 2007. 81(3): p. 559–75.

16. Yang, J., et al., GCTA: a tool for genome-wide complex trait analysis. Am J Hum Genet, 2011. 88(1): p. 76–82.

17. Rousset, F., genepop’007: a complete re-implementation of the genepop software for Windows and Linux. Mol Ecol Resour, 2008. 8(1): p. 103–6.

18. Danecek, P., et al., The variant call format and VCFtools. Bioinformatics, 2011. 27(15): p. 2156–8.

19. Kim, D., et al., TopHat2: accurate alignment of transcriptomes in the presence of insertions, deletions and gene fusions. Genome Biol, 2013. 14(4): p. R36.

20. Trapnell, C., et al., Differential gene and transcript expression analysis of RNA-seq experiments with TopHat and Cufflinks. Nat Protoc, 2012. 7(3): p. 562–78.

21. Acosta, J.R., et al., Single cell transcriptomics suggest that human adipocyte progenitor cells constitute a homogeneous cell population. Stem Cell Res Ther, 2017. 8(1): p. 250.

22. Hilgendorf, K.I., et al., Omega-3 Fatty Acids Activate Ciliary FFAR4 to Control Adipogenesis. Cell, 2019. 179(6): p. 1289–1305 e21.

23. Koza, R.A., et al., Changes in gene expression foreshadow diet-induced obesity in genetically identical mice. PLoS Genet, 2006. 2(5): p. e81.

24. Lee, Y.H., et al., Microarray profiling of isolated abdominal subcutaneous adipocytes from obese vs non-obese Pima Indians: increased expression of inflammation-related genes. Diabetologia, 2005. 48(9): p. 1776–83.

25. LaRochelle, W.J., et al., PDGF-D, a new protease-activated growth factor. Nat Cell Biol, 2001. 3(5): p. 517–21.

26. Dani, C. and A. Pfeifer, The complexity of PDGFR signaling: regulation of adipose progenitor maintenance and adipocyte-myofibroblast transition. Stem Cell Investig, 2017. 4: p. 28.

27. Nishimura, S., et al., ENPP2 contributes to adipose tissue expansion and insulin resistance in diet-induced obesity. Diabetes, 2014. 63(12): p. 4154–64.

28. Zhang, J., et al., Differential Expression of Cell Cycle Regulators During Hyperplastic and Hypertrophic Growth of Broiler Subcutaneous Adipose Tissue. Lipids, 2015. 50(10): p. 965–76.

29. Wei, P., et al., RNF34 is a cold-regulated E3 ubiquitin ligase for PGC-1alpha and modulates brown fat cell metabolism. Mol Cell Biol, 2012. 32(2): p. 266–75.

30. Inagaki, T., et al., The FBXL10/KDM2B scaffolding protein associates with novel polycomb repressive complex-1 to regulate adipogenesis. J Biol Chem, 2015. 290(7): p. 4163–77.

31. Chen, Y.Y., et al., Methyl cinnamate inhibits adipocyte differentiation via activation of the CaMKK2-AMPK pathway in 3T3-L1 preadipocytes. J Agric Food Chem, 2012. 60(4): p. 955–63.

32. Lu, S., et al., Role of extrathyroidal TSHR expression in adipocyte differentiation and its association with obesity. Lipids Health Dis, 2012. 11: p. 17.

33. Korsic, M., et al., Gene expression in visceral and subcutaneous adipose tissue in overweight women. Front Biosci (Elite Ed), 2012. 4: p. 2734–44.

34. Gregoire, F.M., C.M. Smas, and H.S. Sul, Understanding adipocyte differentiation. Physiol Rev, 1998. 78(3): p. 783–809.

35. Liu, Z., et al., Genome-wide detection of selection signatures of distinct tail types in sheep populations. Acta Veterinaria et Zootechnica Sinica, 2014. 46, 1721–1732. (In Chinese)

36. Fan, H., Transcriptomic difference analysis for tail adipose tissue of Hulun Buir sheep. Gansu Agricultural University. 2015. (Ph.D. thesis in Chinese)

37. Petrus, P., et al., Transforming Growth Factor-beta3 Regulates Adipocyte Number in subcutaneous White Adipose Tissue. Cell Rep, 2018. 25(3): p. 551–560 e5.

38. Berulava, T. and B. Horsthemke, The obesity-associated SNPs in intron 1 of the FTO gene affect primary transcript levels. Eur J Hum Genet, 2010. 18(9): p. 1054–6.

39. Tokuhiro, S., et al., An intronic SNP in a RUNX1 binding site of SLC22A4, encoding an organic cation transporter, is associated with rheumatoid arthritis. Nat Genet, 2003. 35(4): p. 341–8.

